# An interactive retrieval system for clinical trial studies with context-dependent protocol elements

**DOI:** 10.1101/814996

**Authors:** Junseok Park, Kwangmin Kim, Seongkuk Park, Woochang Hwang, Sunyong Yoo, Gwan-su Yi, Doheon Lee

**Author notes:** Corresponding author: Doheon Lee.

## Abstract

A clinical trial protocol defines the procedures that should be performed during a clinical trial. Every clinical trial begins with the design of its protocol. While designing the protocol, most researchers refer to electronic databases and extract protocol elements using a keyword search. However, state-of-the-art retrieval systems only offer text-based searches for user-entered keywords. In this study, we present an interactive retrieval system with a context-dependent and protocol-element-selection function for successfully designing a clinical trial protocol. To do this, we first introduce a database for a protocol retrieval system constructed from individual protocol data extracted from 184,634 clinical trials and 13,210 frame structures of clinical trial protocols. The database contains various semantic information that enables the protocols to be filtered during the search operation. Based on the database, we developed a web application called the clinical trial protocol database system (CLIPS; available at https://corus.kaist.edu/clips), which enables an interactive search by utilizing protocol elements. CLIPS provides the options to select the next element according to the previous element in the form of a connected tree, thus enabling an interactive search for combinations of protocol elements. The validation results show that our method achieves better performance than existing databases in predicting phenotypic features.

## Introduction

Clinical trial protocols play a primary role in clinical trials [1, 2]. Well-established protocols simplify clinical procedures, help avoid unnecessary protocol amendments, and facilitate preliminary assessments of latent issues. Thus, well-established and optimized protocols contribute to the success of clinical trials not only by reducing costs but also by improving efficiency [3, 4].

In recent years, optimized protocol design has become increasingly important with the increase in the cost of clinical trials and the complexity of protocols. According to an investigation conducted in 2019, the drug development cost has skyrocketed in past decades, with the total capitalized cost per approved new drug reaching between $200 million and $2.9 billion [5]. In addition, Grand View Research, a market-research company, recently reported that the market size of clinical trials for testing new medical therapies is estimated to reach $68.9 billion a year by 2026 [6]. At the same time, they showed the capitalized clinical trial cost has grown by compound annual growth rate of 5.7% during the forecast period. The highly sophisticated nature of modern drug development is the cause of this upward trend in protocol complexity and work load.

Despite the importance of clinical trial protocols and the increasing demand for streamlining them, contemporary development of clinical trial protocols is rather inefficient. The Tufts Center for the Study of Drug Development (Tufts CSDD) reported that 57% of analyzed protocols had at least one major amendment, and 45% of these amendments had “avoidable” reasons that originated from imperfect protocol design, such as design flaws or recruitment difficulties [7]. Moreover, amendments and changes of clinical trial protocols were concentrated in phase III of the trials, resulting in greater impact costs and implementation efficiency [8, 9].

Because there is a clear demand for a better method of protocol design, there have been various attempts to aid the protocol design process [10, 11]. These approaches can be categorized into two groups: expert guidelines and computerized systems. Computerized systems can be further subdivided into automated and database systems.

The advantage of using a computerized system is that they are quick and allow iterated searches for previous protocols related to the study of interest. In this regard, computerized systems are based on information retrieval technology.

First, existing expert guidelines help researchers design their own trial protocols [12, 13]. While referring to expert opinions guarantees the credibility of the protocol design, two obvious limitations are that the protocol can only be applied if credible guidelines exist in that particular clinical field and not all guidelines offer specific values for all elements of trial design. Consequently, the determination of these specific elements relies on the subjective intuition of individual researchers.

Second, computerized system approaches offer an automated method for designing a protocol. For example, a context-aware architecture for clinical trial protocol design composed of a decision support module and semantic search engine has been developed [14]. Although an idea of constructing an automated system was an innovative approach is proposed, this system offers only limited performance. For instance, the idea focuses on creating scientific queries for finding information about a clinical trial protocol and retrieving only related papers through queries. Furthermore, the web-based service is no longer available.

Current state-of-the-are computerized systems use a database of clinical trial protocols. These databases contain extensive information on previous clinical trials and clinical experiences covering a wide range of clinical fields [15-17]. Researchers can retrieve information from specific clinical trials according to their purpose. However, current clinical trial databases offer only limited support in searching for clinical trial protocols. The clinical trial database system uses medical subject heading (MeSH) terms but does not cover all of the text in the protocols, preventing a truly semantic search [15]. Moreover, the current databases do not allow structural protocol searches to retrieve context-dependent protocol elements. A biomedical literature database system (PubMed) could be used to search for clinical trial protocols [18]. However, this would not be efficient because additional work would be required to extract the necessary information from the retrieved literature.

To overcome the limitations of the current database systems, we present clinical trial protocol database system (CLIPS) that enables a semantic search for the core contents of clinical trial protocols, along with filterable semantic features and frame structures for the protocols. Furthermore, our system efficiently finds clinical trial protocols using a query refinement method. To resolve the difficulty of retrieving specific protocols from a database of complex structures, we developed a graph-based querying system (Fig 1).

**Fig 1. Overview of CLIPS development process.**

## Definitions

Clinical trial protocols consist of several elements that can be grouped into factors according to their characteristics [10]. We define the terms “element” and “factor” as follows.

- Element: individual items constituting the clinical trial protocol. An element has a value that defines the protocol. For example, “model” is an element, and the value of this element can be “crossover.”
- Factor: a common characteristic of grouped elements. A factor can have multiple elements. For instance, “model” and “allocation” elements are used to design a protocol. Thus, they belong to the “design” factor. Another example is the “enrollment type” and “gender” elements which determine the subject of a protocol and are part of the “subject” factor.

## Related work

### Guidelines

The retrieval of documents containing the contents of a protocol design is one method of determining a clinical research protocol. The document containing guidelines covers the overall information of clinical research protocols. This is the most basic approach for gathering information to develop a protocol. Chan et al. proposed SPIRIT, which is a high-quality guideline containing 33 checklist items for the development of a clinical trial protocol [1]. Meeker-O’Connell et al. developed a principle document that defines the factors needed to assure patient safety and reliability in a trial [19]. Moreover, some guidelines specify protocols for examining the efficacy of food or food components for specific diseases [20]. For instance, documents comprising gut health and immunity, diet-related cancer, and atherosclerosis are included [21-23]. However, guideline-based approaches use subjective judgement in determining what information is included [24]. This limitation can result in different outcomes depending on the user.

### Database systems

Database-based information retrieval systems can be utilized to retrieve clinical protocols. Zarin et al. developed clinicaltrials.gov, the largest database-based retrieval system, by collecting all published clinical trial documents, including regulatory mandates and a broad group of trial sponsors [17]. Tasneem et al. established and operated a relational database containing all clinical trials registered with clinicaltrials.gov [15]. Furthermore, systems for protocol retrieval use general document retrieval technologies, e.g., PubMed, Scopus, Web of Science, and Google Scholar [25-28]. However, current database-based retrieval systems have limited ability for protocol-specific search objectives, such as retrieving the protocol structure or selecting context-dependent protocol elements sequentially.

### Intelligent systems

Intelligent systems are an effective approach for retrieving clinical trial protocols. Tsatsaronis et al. developed an intelligent system based on a context-aware approach for automated protocol design [14]. Their system supports study- and domain-driven searches. Study-driven searches use the parameters (i.e., condition, intervention) of a particular trial as provided by a researcher. In domain-driven searches, a researcher selects options according to the study domain, and then the system automatically searches and categorizes the retrieved information. However, such a system is currently not accessible. We assume that the author terminated the system.

## Methods

We developed a clinical trial protocol database system. To accomplish the objective, we proposed step-by-step methods including database development, semantic feature generation, and web-based retrieval system. We constructed the database from a public database of clinical trials and organized essential data to reflect the structure of protocols. Semantic feature generation is a core part of the clinical trial protocol retrieval system. We generated filterable semantic features to offer context-specific search for the protocols from the original text based on named entity recognition tools. The semantic features consist of phenotype, gene, and chemical compound. Finally, we made a web-based protocol retrieval application. Text-based search cannot scrutinize the complex structure of the protocols. Therefore, we devised a graph-based search interface as a query refinement method.

### Database development for a protocol retrieval system

A clinical trial protocol presents the structure of a clinical trial and is composed of various elements that can be clustered into key factors. In this study, we defined five key factors based on a previous baseline research[10, 11]: design, subject, variables, statistical issues, and descriptions. The design factor determines how the trial is structured and modelled to measure data generated during the trial. The subject factor determines who is eligible to participate in the trial and how they are treated to ensure generalizability of the target population. The variables are the parameters to be measured to evaluate the efficacy or safety of a drug or treatment. The statistical issues describe how the clinical trial will be analyzed, specifying sampling procedures or statistical significance. Finally, the description factor covers additional information such as the organization, different phases, and additional explanations of the protocol or trial itself.

We selected and clustered elements from Aggregate Analysis of ClinicalTrials.gov (AACT), which was released on March 27, 2015 [15]. To do this, we first downloaded a dump file of the AACT database and completely overhauled the loaded database to give 42 tables of 270 columns. Then, we classified the data types into four elements: categorical, value, description, and not union (N/U). Categorical-type elements contain categorical variables, and the sequential selection of these elements can determine the protocol structure (S1 Data). Value-type elements include interval and ratio data, which are important values in key factors. Description-type elements contain additional explanatory text, numeric values, and abbreviated words or dates for the description factor. The N/U-type element consists of primary keys, foreign keys, and database management values.

We clustered the categorical and value types into the pre-defined five key factors according to the above mentioned criteria, including design, subject, variable, and statistical issue factors (Table 1).

**Table 1.**
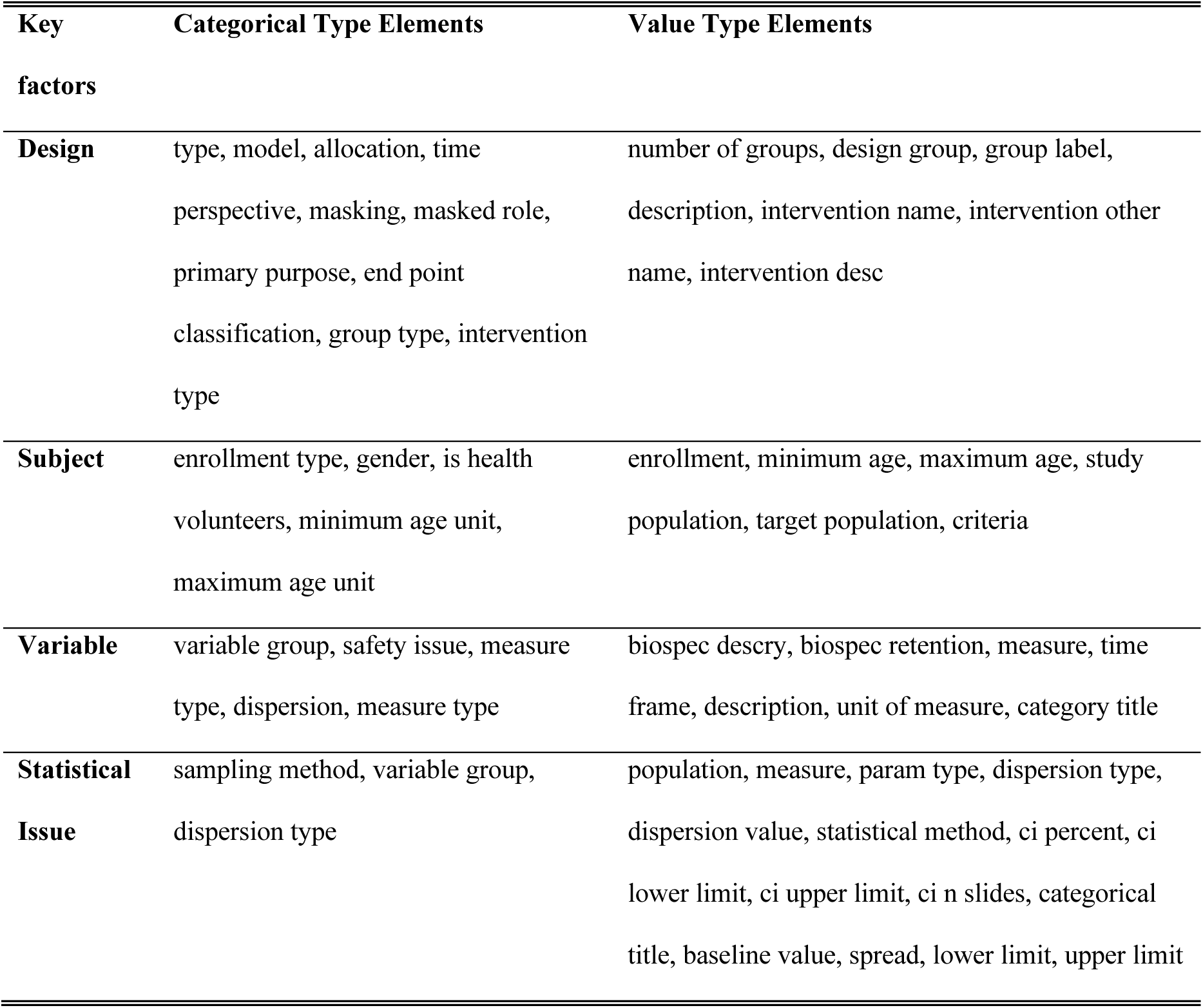
Key factors and their elements in CLIPS.

We designed a table schema and amassed the data compilation progress. The N/U elements were eliminated because we constructed a relational table with key-value attributes, discarding unnecessary keys and values for database management. The use of key-value attributes makes it possible to search the skeleton structure of a clinical trial protocol efficiently and effectively manage the data storage for deploying inconsistent data [29]. We designed a table schema accounting for these aspects (Table 2). The next step was data compilation. We organized the element values by resolving typographical errors and reflecting dependency structures. Then, we removed some control characters and type conversions. Furthermore, we removed ambiguous design types, which are null, and expanded the access and observations (patient registry) to search for specific clinical trial protocols. As a result, we collected 184,634 clinical trial protocols and their detailed information. The resulting database can be used to optimize query refinement for retrieving protocol information.

**Table 2.** Table schema for clinical trial protocol.

| Column Name | Column Type |
| --- | --- |
| ClinicalTrialID | Char(10) |
| Design | JSON |
| Subject | JSON |
| Variable | JSON |
| Statistical Issue | JSON |
| Description | JSON |

### Semantic filtering feature generation

Although we developed a clinical trial protocol database by using a frame structure so that all protocols had a similar structure, the similarity of frame structures does not guarantee similarity of the detailed protocol content. MeSH offers a potential solution and is used for indexing and cataloging clinical trials in ClinicalTrials.gov and AACT [15, 20]. However, MeSH has limited coverage that does not extend across the spectrum of various biomedical terminologies [30]. To solve this limitation, various biomedical semantic features were extracted to find or filter similar clinical trials in the searched structure.

We generated filterable semantic features related to the conditions and interventions that are considered significant in clinical trials, resulting in a subdivided semantic similarity search. The condition was a phenotype, including any diseases and disorders, observed during clinical trials as well as reported symptoms. The disease-specific phenotype is a set of observable characteristics. Drugs commonly refer to intervention which is the focus of a clinical trial, and they can involve chemical compounds [31]. Similar clinical trials can be searched for or filtered through each of the corresponding elements. In addition, the identification of similar target genes or proteins is a promising method of searching for similarities among chemical compounds and phenotypes, as they are a molecular proxy that links them [32, 33]. Thus, we applied named entity recognition (NER) to the phenotypes, chemical compounds, and genes to enable a semantic search, for which Semantic filters were employed for the following description elements: brief title, official title, brief summary, detailed description, keywords, and conditions (Fig 2).

**Fig 2. Generation of data for semantic search.**

### Phenotype

We extracted semantic features to represent disease specific phenotype words. The unified medical language system (UMLS) is a repository of integrated biomedical terminologies, and thus we used UMLS2015AB to process phenotype words [30]. To employ NER on descriptive values, we applied Metamap 2016 and cTakes 3.2.2 [34, 35]. We combined each result and removed duplicates using the above-mentioned tools to synthesize the advantages [36]. Next, we selected 15 semantic types, which are considered disease phenotypic types, and removed the other types from the results (Table 3). As a result, disease phenotypic features with unique concept IDs were generated for each clinical trial.

**Table 3.** Selected phenotypic types from UMLS.

| Entity Type ID (TUI) | Entity Type Name |
| --- | --- |
| T038 | Biologic Function |
| T039 | Physiologic Function |
| T041 | Mental Process |
| T019 | Congenital Abnormality |
| T020 | Acquired Abnormality |
| T033 | Finding |
| T034 | Laboratory or Test Result |
| T046 | Pathologic Function |
| T047 | Disease or Syndrome |
| T048 | Mental or Behavioral Dysfunction |
| T049 | Cell or Molecular Dysfunction |
| T184 | Sign or Symptom |
| T190 | Anatomical Abnormality |
| T191 | Neoplastic Process |
| T037 | Injury or Poisoning |

### Chemical compound

We applied NER to chemical compound entities from the descriptions given by ChemSpot [37]. ChemSpot provides Chemical Abstract Service (CAS) IDs and International Chemical Identifiers (InChI) but does not provide standard InChIKeys. The InChIKey is the compacted version of InChI, and the standard InChIKey is a stable identifier for reflecting the identifier version designation [38]. Moreover, standard InChIKeys are considered to provide equivalent descriptions between compounds in drug discovery [39]. To take advantage of the standard InChIKeys for chemical compound entities, we examined the original words of the NER-processed entities by using ChemSpider [40]. The simple application programming interface (API) of ChemSpider was exploited, allowing us to generate chemical compound entities with standard InChIKeys, InChIs, and the simplified molecular-input line-entry system notation.

### Gene

We appended gene entities in the elements of semantic filters. The gene annotation tool, Moara, was used for gene NER, considering Moara is capable of performing both recognition and normalization of gene entities, recognizing entities and their positions in the input text, and linking the entities to gene IDs in a known gene database [41]. Moara provides various preconstructed machine learning-based models for various organism species. For our task, we adopted the human-oriented model. For gene normalization, we obtained lists of gene IDs corresponding to each gene entity. The gene ID with the highest score was selected and mapped to the recognized gene term.

### Web application development for query refinement

Clinical trial protocols have increasingly complex structures [42]. The complexity of protocol level is inversely related to clinical trial performance, as complex protocols negatively impact factors such as protocol amendment rates, patient recruitment, and retention rates [43]. In addition, the increasing complexity of the protocols hinders the design of new protocols, because clinicians referring to previous clinical trials to design a protocol inevitably face difficulties in searching for suitable examples. Thus, from a query refinement standpoint, we developed the CLIPS web application to provide a graph querying interface for retrieving information about reliable clinical trial protocols, rather than a text querying interface that cannot visualize the dependency among prior elements affecting the protocol structure [44, 45].

We defined categorical-type elements as the frame structures of protocols. Although we have provided default orders of the elements, the user is free to choose the order. Once a decision about the order has been made, the user can find varying combinations of protocol elements via the graph-based search interface. Dependent elements are retrieved from the database in real-time, and the user can confirm the number of existing protocols corresponding to selected elements. The user also can search for other protocol frame structures after clipping the selected structure to the user clip pane. To search for complete information about the selected protocol structures, the user must click on a clipped protocol to examine the details of the selected protocol and trial information. Furthermore, the user can add semantic filters when searching for complete information to reduce or focus on certain areas of clinical trials. The data flow of the interface is illustrated in Fig 3.

**Fig 3. Example of background data-flow on CLIPS from a research question in a clinical trial.**

The backend of the interface was developed using Node.js [46] and the visualization of the interface was manipulated by d3.js [47]. We developed custom functions on d3.js to show each element title and protocol count for the user’s selection. To implement semantic filtering of the user’s free-text input, a backend engine is connected to the representational state transfer (REST) NER API. This provides NER of processed entities and types, allowing relevant entities to be searched for in the database. We designed the system architecture to combine the interface application, APIs, and database for stable operation in a cloud-computing environment (Fig S1).

## Results

### Database

We collected 184,634 clinical trial protocols and 13,210 frame structures of clinical trial protocols and extracted 5,765,054 phenotypes, 1,151,053 chemical compounds, and 222,966 gene features for semantic filtering (Table 4). Furthermore, we designed a continuous process of data update so that the protocol methods could evolve naturally, thus enhancing the quality of the database (S2 Fig). In conclusion, we developed a database system that efficiently retrieves information about existing clinical trial protocols for use in designing new clinical trial protocols.

**Table 4.** Table schema for clinical trial protocol.

| Entity | Rows (Unique) |
| --- | --- |
| Clinical Research Protocol | 184,634 (184,634) |
| Frame Structures | 13,210 (13,210) |
| Phenotype | 5,765,054 (18,438) |
| Chemical Compounds | 1,151,053 (12,792) |
| Genes | 222,966 (4,705) |

### Application

The web application provides a service for retrieving protocol structures and inquiring about protocol information. The user goes through four stages in using this service: (1) setting the order of the protocols; (2) designing the protocol structure by selecting the elements that correspond to each sequence; (3) setting various functions required to search for the desired protocol information; and (4) providing the desired protocol information to explore the contents in detail. We developed the necessary interfaces to perform all of these processes (Fig 4).

**Fig 4. System overview.**

Before retrieving the protocol structure, the user should define the protocol sequence. This process uses a drag-and-drop interface to sort the list into one box and set the order. This allows users to work more intuitively [48]. After determining the protocol sequence, the vector-based collapsible and zoomable tree diagram visualization interface, which is an uncomplicated tool to navigate the protocol structures with selected elements [49, 50]. The loaded protocol data are assembled into a hierarchical data structure. This visualization is rendered as a relation tree with a parent/child structure. The user must click on the edge of the tree to add the next-step protocol. Conversely, to remove protocol edges from the current stage, the user must click on the parent edge of the previous step. The entire data structure is synchronized and updated every time the process occurs [51].

After the protocol structure has been retrieved, the user obtains protocol information based on the selected protocol structure. We developed a function called Clip to back up the protocol design. This allows the user to reuse previously selected protocol structures and receive corresponding study information. In addition, a protocol-information-filter function allows the user to retrieve the study information. The user searches for a disease and generates a label that contains the disease code. The protocol information is then filtered according to the set label. The resulting data are rendered as a table, which can be sorted with respect to the column entities to focus on and export specific data. When exploring detailed protocol information, our system transforms the data into a collapsible interface instead of providing raw text.

Although protocol design concepts have evolved globally, the development of tools to design clinical trial protocols is trivial [52-54]. Our aim was to simplify clinical trial retrieval and the design stage by developing a dedicated interface. We expect this to be the starting point for the creation, sharing, and development of more clinical trial protocols.

## Validation

### Technical validation

The goal of CLIPS was to provide an information retrieval method that can search complex clinical trial protocols. For this, we developed a search tool that can build and utilize a database suitable for the protocol structure. Furthermore, we created semantic features in CLIPS using text-mining methods. As a result, it was possible to perform accurate searches using the contextual meaning of the protocol. To evaluate the performance of CLIPS, we attempted to verify whether the semantic filter results in better performance than a keyword search.

For the technical validation of the semantic filter of CLIPS, we used the relation information between clinical trial protocols and corresponding disease conditions as collected from clinicaltrials.gov[17]. As this disease condition assignment is manually curated by experts and does not originate from the protocol itself, it can be used as a gold standard to evaluate the semantic filters of CLIPS.

The gold standard set of disease conditions and corresponding trial protocols was obtained by crawling the topic page of clinicaltrials.gov[17]. Among 25 conditional categories provided by clinicaltrials.gov, the “Cancers and Other Neoplasms” condition category was selected because it covers 44.74% of the total protocol set. Consequently, the corresponding trial protocols and corresponding disease conditions were identified. For instance, in our gold standard set, 353 distinct protocols were associated with the disease condition “Abdominal neoplasm.” As a result, a set of 82,584 distinct protocols corresponding to 520 disease conditions was compiled (as of July 12, 2017) and used as a gold standard set for technical validation.

The semantic search performance of CLIPS was validated using the following procedure. In CLIPS, search keywords containing the disease condition names were supplied as input queries to the system. The semantic entities were translated from the search keyword through the text-mining-based models described in the previous section. The results were obtained by conducting a search using the translated semantic entities from the CLIPS database. Exact matching with the AACT database was used as a baseline. CLIPS and AACT database were configured on a single local server.

We used a condition name (e.g., Adrenocortical Carcinoma) as a search keyword to retrieve the condition field of the source database and semantic entities (e.g., C0206686) from CLIPS. We then validated our retrievals by comparing the number of retrieved identifiers (NCTID) of protocols (S2 file)[55] and calculated the precision, recall, and F1-score of the results. Precision refers to how accurately the model is categorized and is calculated as the ratio of properly categorized data to the total data (1). Recall is the number of positively classified data divided by the original number of positive data (2). F1-Score is a harmonic mean considering the complementary characteristics of precision and recall (3), and it is commonly used to compare the performance of different information retrieval systems [56]. True positive + false positive is the count of all NCTIDs retrieved from each database. True positive is the count of the retrieved NCTIDs of each database intersected with the gold standard. True positive+ false negative is the total number of NCTIDs in the gold standard corresponding to each input condition name.

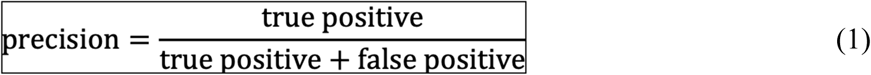

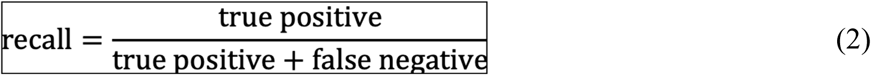

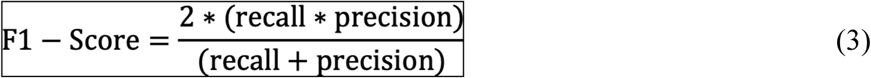

The F1-Score of CLIPS (0.515) was higher than that of the keyword search (0.38) (Fig 5). The precision of CLIPS was 0.437, which was slightly lower than that of the keyword search (0.668), but it outperformed the keyword search by more than a factor of two in terms of recall (0.63 and 0.26, respectively). As higher recall values are a positive factor in a clinical trial design, CLIPS can retrieve more protocols that provide more suitable references for protocol design.

**Fig 5. Evaluation results of keyword search and using semantic filter of CLIPS for “Cancer and Other Neoplasms” categorized conditions.**

### User experience

As described earlier, the CLIPS search system was developed for different purposes than those of the existing clinical trial search systems. CLIPS is intended to assist in the effective design of a specific trial protocol, whereas existing search systems are generally used to process various information on clinical trials. Therefore, to evaluate the performance of CLIPS, an evaluation method that reflects this purpose should be constructed. As the ultimate goal of a search system is to help users collect the information they require, the subjective satisfaction of users is a significant measure of performance. Thus, the evaluation of a search system should be able to quantify the subjective impressions of users as well as objective indicators. By considering these factors, we conducted an evaluation trial that compared CLIPS with the conventional search system provided by clinicaltrials.gov.

Ten participants aged between 24 and 33 were recruited from a group of experts in the field of bioinformatics (Bio and Brain Engineering Department of Korea Advanced Institute of Science and Technology, Republic of Korea) They included two undergraduates, three master’s students, four PhD candidates, and one postdoctoral researcher. All participants provided informed consent before conducting the evaluation trial. The participants were assigned the task of finding a suitable clinical trial set for a simulated problem. Two separate tasks were given to the participants, who were asked to construct the most common previous protocol design under the given trial conditions and research questions and perform each task using each of the two search systems within the time limit (5 min each). To construct the most common trial design, participants had to collect information about the various elements of the trial protocol (Fig 6). After the task, participants completed a questionnaire on their subjective satisfaction regarding the system. The questionnaire consisted of six questions that enquired about the participants’ satisfaction using a 7-point Likert scale [57].

**Fig 6. Tasks given to participants for evaluation.**

Participants were observed to perform better when using CLIPS than when using the clinicaltrials.gov search system. By using CLIPS, participants obtained more answers within the time limit, and the average time required to perform the task was shorter. The number of clinical trials retrieved from CLIPS was less than that from clinicaltrials.gov because it was possible to apply more detailed search filters to narrow the search scope. Participants were more satisfied with CLIPS than the existing search systems, as evidenced by the average score of the questionnaire responses (Table 5). These results show that CLIPS can be effectively used to retrieve certain types of trial protocols.

**Table 5.**
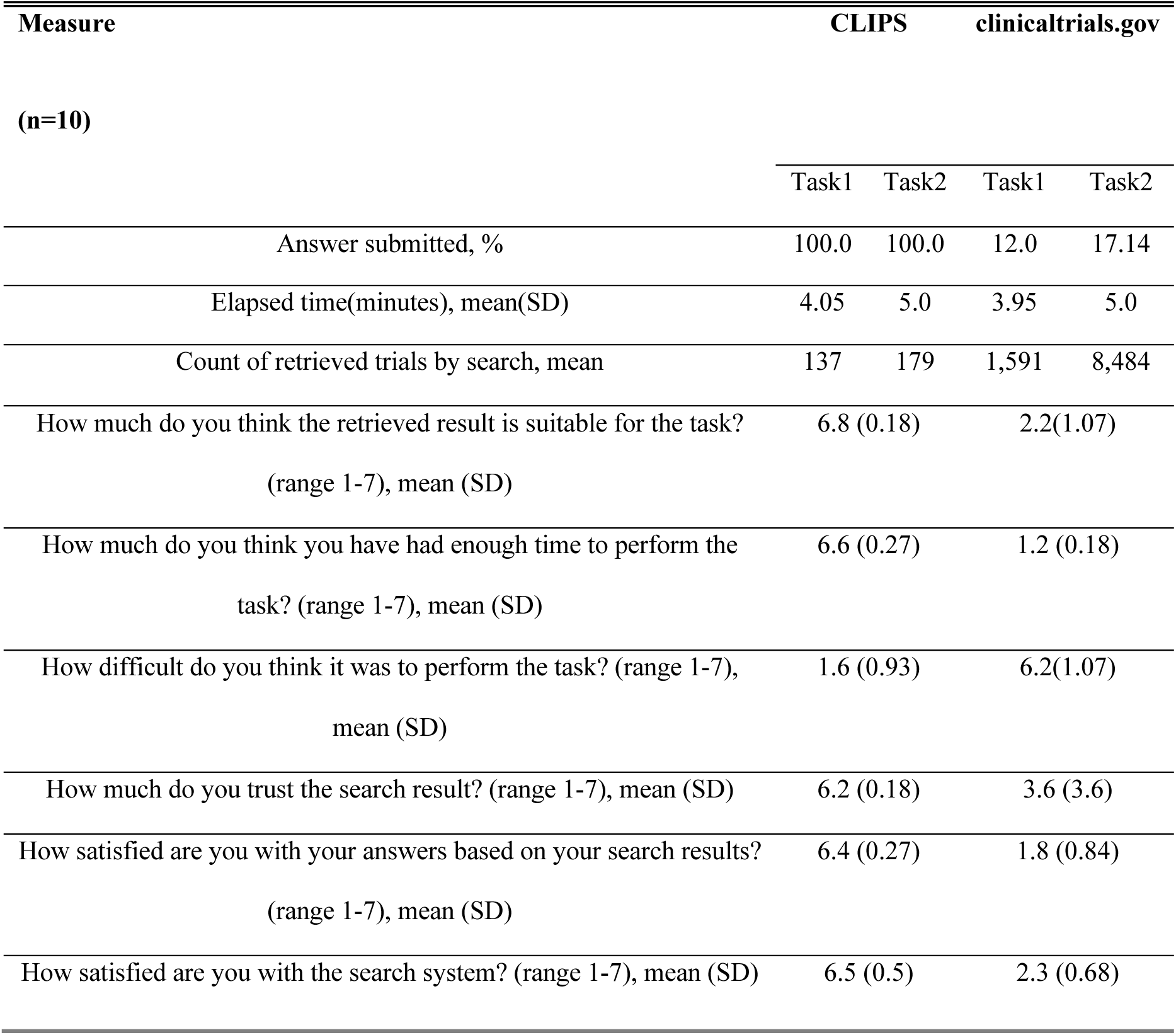
Evaluation results.

| Measure | CLIPS |  | clinicaltrials.gov |  |
| --- | --- | --- | --- | --- |
| (n=10) | Task1 | Task2 | Task1 | Task2 |
| Answer submitted, % | 100.0 | 100.0 | 12.0 | 17.14 |
| Elapsed time(minutes), mean(SD) | 4.05 | 5.0 | 3.95 | 5.0 |
| Count of retrieved trials by search, mean | 137 | 179 | 1,591 | 8,484 |
| How much do you think the retrieved result is suitable for the task?<br>(range 1-7), mean (SD) | 6.8 (0.18) |  | 2.2(1.07) |  |
| How much do you think you have had enough time to perform the<br>task? (range 1-7), mean (SD) | 6.6 (0.27) |  | 1.2 (0.18) |  |
| How difficult do you think it was to perform the task? (range 1-7),<br>mean (SD) | 1.6 (0.93) |  | 6.2(1.07) |  |
| How much do you trust the search result? (range 1-7), mean (SD) | 6.2 (0.18) |  | 3.6 (3.6) |  |
| How satisfied are you with your answers based on your search results?<br>(range 1-7), mean (SD) | 6.4 (0.27) |  | 1.8 (0.84) |  |
| How satisfied are you with the search system? (range 1-7), mean (SD) | 6.5 (0.5) |  | 2.3 (0.68) |  |

## Discussion

Clinical trial protocols are the foundation for planning, approving, conducting, and reporting clinical trials [1]. They include general information, objectives, trial design, the selection and withdrawal of subjects, treatment, safety assessments, quality control procedures, and record keeping processes [58]. This study aimed to develop an efficient system of providing the information necessary for clinical trial protocol development. In particular, we have made it possible to find previous protocols of the desired type using the structural features of the protocol composed of context-dependent protocol elements. Furthermore, semantic filtering was included to ensure the retrieval of relevant protocol context information.

CLIPS can search for protocols or specific disease names and structures. In addition, CLIPS can be used for a combination of structural searches, structural order searches, semantic searches, or searches including both structure and semantic context. For instance, our system can perform the following functions:

We suppose that a user has a plan to develop a clinical trial protocol about Cardiovascular Infections, and the user will perform an observational study and should decide the sampling method.

1. Input ‘Cardiovascular infections’ to the semantic filtering input box.
2. Ordering elements as follows: 1. Type, 2. Sampling method. The system orders the rest of the elements automatically.
3. Click ‘complete’ on the protocol order interface.
4. Select ‘node’ on the graph-based search interface.
5. Click ‘type’ and then wait until the next element result is provided.
6. Click ‘observational’, and then, the user can find ‘Non-probability sample’ is the sampling method to the specific disease.
7. Click ‘Clips’ below the graph-based search interface
8. Click clipped protocol structure, and then the system provides detail information of 6 searched protocols.
9. The user can follow or download more detailed protocol information based on a detailed description of the searched protocols from explanation interface such as model, enrollment type, time perspective and outcome variables.

Similar to the above results, we believe that many clinicians will be able to utilize our system to design more reliable clinical trial protocols. The developed semantic filter can be used to search for protocols and can be used for drug discovery using the retrieved protocols. CLIPS provides search results as a downloadable file containing the semantic filters as well as protocol structure information. This ensures the wide coverage of the protocol search. For example, we used UMLS as a phenotype semantic filter. UMLS is an integrated terminology system that combines biomedical terminologies including SNOMED-CT, MeSH, and MedDRA [59]. Clinical terms in SNOMED CT have been integrated into the UMLS metathesaurus since 2003 [60]. For instance, coverage of the HPO term is higher than in SNOMED CT [61]. According to Bodenreider’s study, UMLS covered 54% of the HPO phenotype terms, whereas SNOMED CT covered only 30% [30]. Based on the semantic filter, researchers can screen chemical compounds of drug candidate substances, regardless of whether they are known to be effective, from the previous retrieved protocols [62]. Gene and phenotype can also be used for drug efficacy screening or efficacious drug combinations in massive biological networks using a similar approach [63-65]. Furthermore, users can download the entire database. The efficacy weight of edges can then be predicted, the predicted pathways validated in large-scale biological networks and be utilized to find their maximum therapeutic benefits [66, 67].

The present study is limited in terms of user-experience validation. This is because clinicaltrials.gov and CLIPS have different objectives. Clinicaltrials.gov was developed to register clinical trials[17]. The registered information can be retrieved by clinical researchers, patients, and families of patients. This is a different objective from that of CLIPS, which is specially designed for the protocol search task. As the objective is different, the method is different. Therefore, it cannot be claimed that the clinicaltrials.gov offers worse performance than CLIPS, although the user-experience validation experiment showed a low score for use of clinicaltrials.gov. The satisfaction score of CLIPS is high in terms of the objective of protocol searching. If clinicaltrials.gov were to include the search function of CLIPS, it would offer comprehensive and specific information search ability to clinical researchers.

## Conclusion

Clinical trial protocols are a crucial factor in allowing clinical trials to achieve their primary purposes. However, clinical researchers tend to design clinical trials according to their individual expertise. This type of bias or ambiguity may lead to inconsistencies and objectivity problems during the designing of clinical trial protocols. To solve these problems, an information retrieval system for clinical trial protocols is needed. In this study, we developed a clinical trial protocol database system and an information retrieval web application. The database contains design, subject, variable, statistical issue, description and structure of clinical trial protocols. The web-based protocol retrieval system provides a graph-based search interface based on the structure, and then users can find relevant information of a protocol. Furthermore, the database also includes semantic features to serve the context-specific protocol search. Unlike the previous clinical trial database system, our system has the following two main strengths: (i) provide structural information to present simplified element-wise selection and (ii) extend the search field based on filterable semantic features to do a context-specific search for clinical trial protocols. We believe that CLIPS will be a major resource for clinical trials and will be of interest to clinicians and pharmaceuticals or even regulatory agencies by providing information about clinical trial protocols conveniently. This study has described the formulation of the CLIPS database system and explained its implementation and advantages over existing keyword-base search systems.

The whole database is available for download (http://corus.kaist.edu/clips).

## Acknowledgment

We would like to thank David Sharpe for his cooperation in using ChemSpider APIs and Seyol Yoon for helping us select phenotypic TUIs from UMLS.

## Supporting information

**S1 File. Supplementary Data 1.**

**S2 File. Supplementary Data 2.**

**S3 File. Supplementary File.**

**S1 Fig. CLIPS service architecture on Amazon web service.** Domain name service (DNS) uses the KAIST domain server to use kaist.edu. A user accesses the CLIPS service through the DNS. When accessing CLIPS through the DNS, the interface elastic compute cloud (EC2, https://aws.amazon.com/ec2/) is called, and it displays a screen to the user. Interface EC2 connects to API Engine EC2 to process the data requested by the user. If the user uses a semantic filter, API engine EC2 transfers the input value of the user to text mining EC2, and then it receives the result. Particularly, text mining EC2 is composed of Metamap1, Moara2, and Chemspot3Dockers4, which we customize for our service in the elastic container service (ECS, https://aws.amazon.com/ecs/) group. To provide the data requested by the user, API engine EC2 receives the searched result from the CLIPS relational database service (RDS, https://aws.amazon.com/rds/) in which the clinical trial protocol data are stored, and it transfers the result to interface EC2. Furthermore, we use elastic load balancing (ELB, https://aws.amazon.com/elasticloadbalancing/) for stable service traffic control, and ELB is required to make requests for the EC2 groups that are grouped into the auto scaling group (https://aws.amazon.com/ec2/autoscaling/).

**S2 Fig. Continuous data update flowchart of CLIPS.** The DB Checker, a database change detection function based on the Quartz job scheduler (http://www.quartz-scheduler.org) built in the CLIPS API engine, operates as follows: (1) DB checker detects whether the source database is changed or not. (2) DB Checker detects whether schema of the database is changed or not (3) Schema change (3-1) In case of schema change, the program cannot process it automatically. Therefore, it needs to analyze manually. The DB checker sends an update notification email to the CLIPS developers using the AWS simple notification service (https://aws.amazon.com/sns/). (4) Schema not changed. Only new data are added. (4-1) The modified dataset is stored in the CLIPS temporary data storage table. (4-2) Protocol structure information is extracted from a temporary data table. (4-3) Semantic features are extracted from the texts of the data using the CLIPS text mining engine. (4-4) The result obtained in the previous step is stored in the CLIPS database, and the update is completed.

**S3 Fig. CLIPS example of protocol structure retrieval by the selection of a categorical type element.** A user selects the type as the first element in the categorical type element and then chooses the intervention category. The second categorical type element is of primary purpose, and the diagnostic is decided among the nine categories included in the element.

