## Supplementary material for "An interactive retrieval system for clinical trial studies with context-dependent protocol elements": Graphical Abstract (from another version of this preprint: version 1, 2019-10-23)

### Interactively retrieving clinical trial protocols for successful clinical trial protocol design

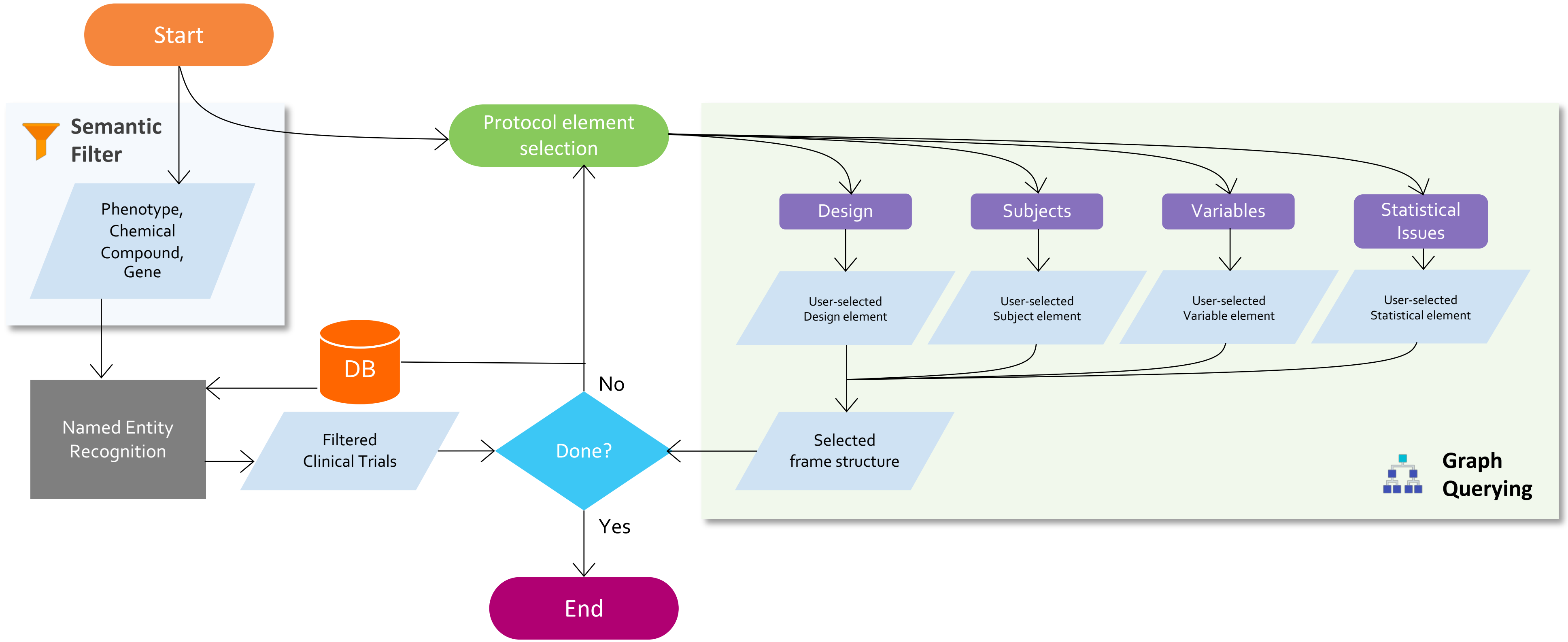
